# Low quality evidence dominates discussion of carbon benefits of alternative grazing strategies

**DOI:** 10.64898/2025.12.09.693242

**Authors:** Jonathan Sanderman, Colleen Partida, Yushu Xia, Jocelyn M. Lavallee, Mark A. Bradford

## Abstract

There is enormous interest in utilizing alternative grazing strategies to improve rangeland condition, increase profitability and decrease the carbon footprint of livestock production via soil organic carbon (SOC) sequestration. Here we present a systematic review and meta-analysis of the literature on alternative grazing strategies and their impact on SOC. Most studies (47 out of 70) failed multiple quality criteria. The 10 studies with 27 observations that met all inclusion criteria showed no change in SOC. Further dividing these observations by study design (controlled small-plot experiment versus observational paired site study) or grazing management style (adaptive or prescribed) or by aridity class or by grassland type failed to reveal any trends. Gains in SOC were only found for a secondary group of 13 studies with 25 observations, primarily composed of paired comparisons where there were unresolved questions about the quality of the site pairings for supporting causal inference. This divergence in results between primary and secondary studies highlights that low-quality evidence dominates the discussion around the climate benefits of alterative grazing strategies, underscoring a critical need for stronger evidence before asserting climate change mitigation benefits from alternative grazing practices.

**For submission to:** Communications Earth & Environment

## Introduction

Grazing of domesticated ruminant animals is the largest land use globally. There are ∼32 million km^2^ of land that cannot support crop production but produce about one third of global protein^1^. Inappropriate management of rangelands has led to large-scale degradation and desertification^2^, resulting in losses of 10’s of billions of tons of soil organic carbon (SOC)^3^.

Reversing rangeland degradation through regenerative grazing practices is being pursued by governments and civil society as a key nature-based climate solution (NbCS), with an estimated potential to sequester 100s - 1000s of millions of tons of CO_2_ annually^4,5^ and exploding interest in paying producers as part of various carbon programs. Yet conflicting evidence of the impacts of grazing management on SOC has caused deep polarization within the scientific and land management communities, and there is a critical need for clarity to drive consensus around this potential NbCS pathway.

Regenerative grazing is a vague term and often refers to a suite of practices including holistic planned grazing, management-intensive grazing, cell grazing, adaptive multi-paddock grazing, time-controlled grazing or short-duration rotational grazing^6^. At the core of these practices, which we call ‘alternative grazing strategies’ in this work, is development of grazing management plans that achieve various production and/or conservation outcomes. These plans can be either prescribed (i.e. grazing schedules are set at the beginning of a season) or adaptive (i.e. grazer movements are based on observations of forage consumption) in nature^7^. Critically, these outcomes are typically considered relative to continuous or season-long extensive grazing strategies, where there is minimal control of livestock movements within large areas. Alternative grazing management plans typically have three key objectives^8^: 1) Improve grass productivity and species composition through adequate rest periods during the growing season; 2) Encourage livestock to consume a broader range of vegetation by increasing stocking density; and 3) More uniformly distribute grazers across a grazed area through fencing and improved water distribution. When these objectives are met, there is evidence of improved floristic and productivity benefits^9,10^ which could, in theory, translate into gains in SOC or at least reductions in soil degradation and associated SOC loss seen in many rangeland ecosystems.

There is a wealth of evidence that optimal grazing intensity (proportion of aboveground net primary productivity consumed by livestock over the course of a year), which for much of the world means lower livestock numbers per unit area, assuming season-long grazing, can increase SOC levels^11^. In contrast, alternative grazing strategies do not decrease grazing intensity per se but instead spread grazing more evenly, meaning results from grazing intensity studies cannot be used to understand the SOC benefits of alternative grazing management. Further, quantitative syntheses of “improved grazing management” largely comprise individual studies that are not equivalent to alternative grazing strategies^12^, questioning the generalizability of the reported management effects. The first meta-analysis^13^ to specifically include a comparison of alternative versus traditional continuous (or season-long) grazing strategies found an ∼28% improvement in SOC under alternative strategies, whereas a more recent meta-analysis focused on Australia^14^ found no discernible SOC effect. Several recent syntheses, including Bai and Cotrufo^4^, cite the values reported that first meta-analysis^13^ as the potential of alternative grazing strategies to increase SOC levels.

Establishing realistic estimates of potential carbon or climate benefits when producers adopt a new livestock management system is important for producers, climate and conservation investors and policymakers. Further, understanding where and under what conditions benefits will accrue is needed for making informed decisions. It has been hypothesized that there might be more of a positive SOC response in more mesic environments where productivity is less constrained by available soil moisture^15^. Following the same logic, there might be more room for carbon accrual in improved pastures because sown grass varieties – often in conjunction with liming, fertilization or irrigation – are more productive than their native counterparts^16^; however, there is growing evidence that diverse native grasslands can outperform sown pastures once climate variability is considered^17^. Finally, whether or not producers respond adaptively to actual forage conditions when scheduling stocking rates and livestock movements can impact SOC outcomes especially if a prescribed rotational grazing management strategy results in over-utilization and degradation of the grassland^7^.

A notable shortcoming of past meta-analyses and syntheses on this topic is that, as is common in meta-analysis^18,19^, the quality of evidence in studies with new direct measurements is not rigorously considered. Yet, it is well established that meta-analyses will result in misleading or uninterpretable conclusions if all data are included without a critical evaluation of the validity of the individual studies^18,20^. Building confidence in the accuracy of the estimated treatment effect – in this case, the effect of alternative grazing management – requires a balance of evidence between highly controlled, well-replicated interventional studies (high internal validity but often conducted at a small scale) and well-designed observational studies conducted at the scale of real-world grazing operations (high external validity). This study evaluates whether such a balance of evidence is available by conducting a systematic review of the peer-reviewed literature where the quality and power of the underlying studies are considered during the data compilation and analysis stages of a traditional meta-analysis. We primarily consider the results based on study quality and experimental design, and interpret these findings using a causal inference lens^19,21,22^. We also use the meta-analysis results to test hypotheses around climate (dryland or non-dryland), grassland type (native or improved) and alternative grazing management style (adaptive or prescribed).

## Results

Our systematic review identified 70 research studies that compared new direct measurements of SOC outcomes for conventional (continuous or season-long) versus alternative grazing strategies. Only 10 studies (termed primary hereafter) met all inclusion criteria (detailed in Table 1), whereas many failed multiple criteria (Figure 1). If a study met all inclusion criteria except the question of whether the control was representative of the pre-intervention management in the treatment plot, it was included in a secondary category of studies (n = 13). Forty-seven studies were excluded due to failing multiple quality criteria. The final dataset for further analysis consisted of 27 comparisons of SOC stocks from 10 primary quality studies and 25 comparisons from the 13 secondary quality studies. Egger’s regression^23^ suggested that there was no publication bias across all studies nor in the primary or secondary study classes respectively (see Figure S1).

**Figure 1.**
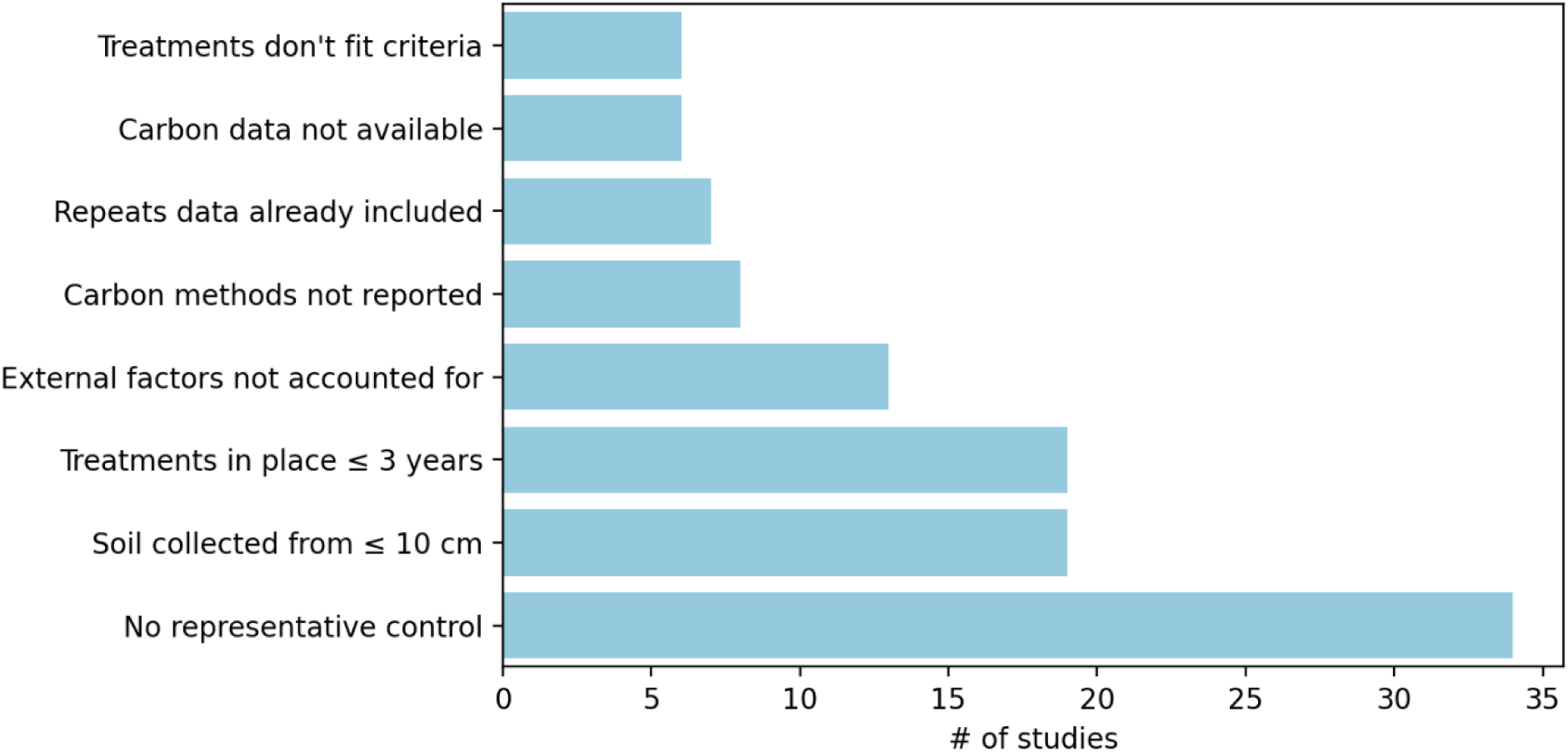
Counts of reasons for individual studies omitted from primary analysis of grazing management impacts on SOC change. Lack of representative control was the most common exclusion criteria. Note: an individual study could fail multiple criteria.

**Table 1.** Inclusion or exclusion criteria.

| CRITERIA | OMIT | INCLUDE AS PRIMARY |
| --- | --- | --- |
| Is the carbon data available in a table or figure, or as a supplemental dataset? | - No | - Yes |
| What methods were used to determine carbon content? | - Missing methodological detail | - Uses loss on ignition (LOI)<br>- Uses dry combustion and reported carbonate removal if carbonates may be present<br>- Uses Walkley-Black |
| How long have the treatments been in place? | - Less than 3 years<br>- Cannot be determined | - Greater than or equal to 3 years |
| To what depth was soil collected? | - Depth of 10 cm or less | - Greater than 10-cm depth |
| Are there significant external factors that may affect SOC sequestration between treatments that are NOT accounted for in data analysis? | - Yes, i.e. differing climate, topography, parent material, soil type NOT accounted for in data analysis (raw means reported) | - No, i.e. no significant factors OR factors are accounted for in data analysis (conditional means reported) |
| Is the control representative of the pre-intervention management in the treatment plot? | - No<br>- Cannot be determined | - Yes |

Perhaps given our inclusion criteria of a minimum sampling depth of 10 cm, the average sampling depth for the included observations was 51 cm with little difference between primary and secondary studies. Study duration (or years of divergent management for paired comparisons) averaged 13 years across all studies, with the paired site designs having an average duration of 15 years and the controlled experiments averaging 10 years.

Rather than uncovering small but real effects - a typical strength of meta-analysis - our analysis showed no significant effects of alternative grazing strategies on SOC. When all primary and secondary quality studies were pooled together the mean natural-log of the response ratio (lnRR) of SOC was 0.05 (95% CI: −0.002 – 0.105) and the mean SOC sequestration rate was 0.29 (95% CI: −0.04 – 0.64) tC ha^-1^ yr^-1^ (Figure 2). When the results were narrowed to only primary studies, the mean SOC response to adoption of alternative grazing strategies was indistinguishable from zero (Figure 2). The net positive effect for ‘All’ studies was driven almost entirely by the secondary study category (mean lnRR = 0.097 and mean SOC sequestration rate = 0.50 tC ha^-1^ yr^-1^), dominated by paired-site studies with only post-intervention measurements and questions around the representativeness of the controls used for comparisons. One-way ANOVA results for lnRR (F = 4.053, p = 0.049) and SOC sequestration (F = 4.037, p = 0.051) confirm these differences between categories.

**Figure 2.**
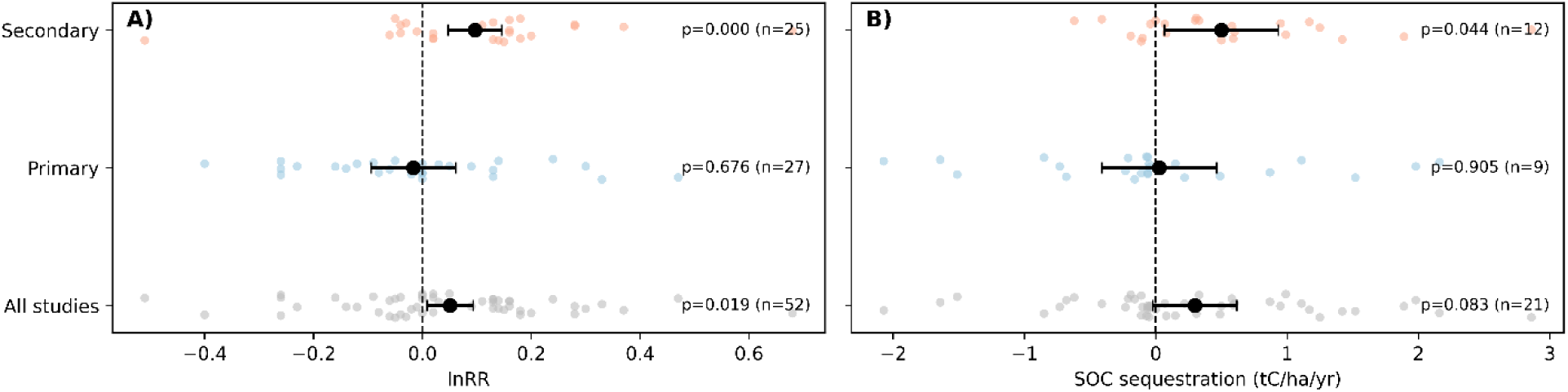
Change in soil organic carbon (SOC) after adoption of alternative grazing management. For the natural-log of the response ratio (lnRR) (A), points show individual effect sizes and black symbols show pooled random-effects meta-analytic estimates with 95% confidence intervals. For SOC sequestration rate (B), points show individual observations and black symbols show study-clustered mean estimates with 95% confidence intervals, calculated from among-study variation because observation-level standard errors were unavailable.

Grouping observations by experimental design also did not uncover any significant responses. The primary studies were split between controlled experiments (n = 11) and paired site observations (n = 16); however, the secondary study category was dominated by paired site observations (n = 20) with only two controlled experimental observations. Overall, controlled experiments had a mean lnRR and mean SOC sequestration rate indistinguishable from zero, while the paired site studies had positive mean lnRR and mean SOC sequestration rates (Figure S4). If the results were narrowed to primary studies only, none of the responses were different from zero (Table S4) suggesting that experimental design alone is not the only factor contributing to the divergence in results between the primary and secondary study categories.

Despite hypotheses suggesting that SOC may respond more readily to grazing in mesic, more productive rangelands ^24^, our findings suggest no difference in SOC response between dryland and non-dryland aridity classes (Figure 3) and no difference between native and improved/sown grasslands (Figure 4). However, when only focusing on the secondary studies, the improved grasslands, which represent all of the non-dryland observations, had a mean lnRR significantly greater than zero (mean lnRR = 0.15, p = 0. 007) while in native grasslands, all in dryland climate zones, there was too much dispersion in the data to detect a significant difference.

**Figure 3.**
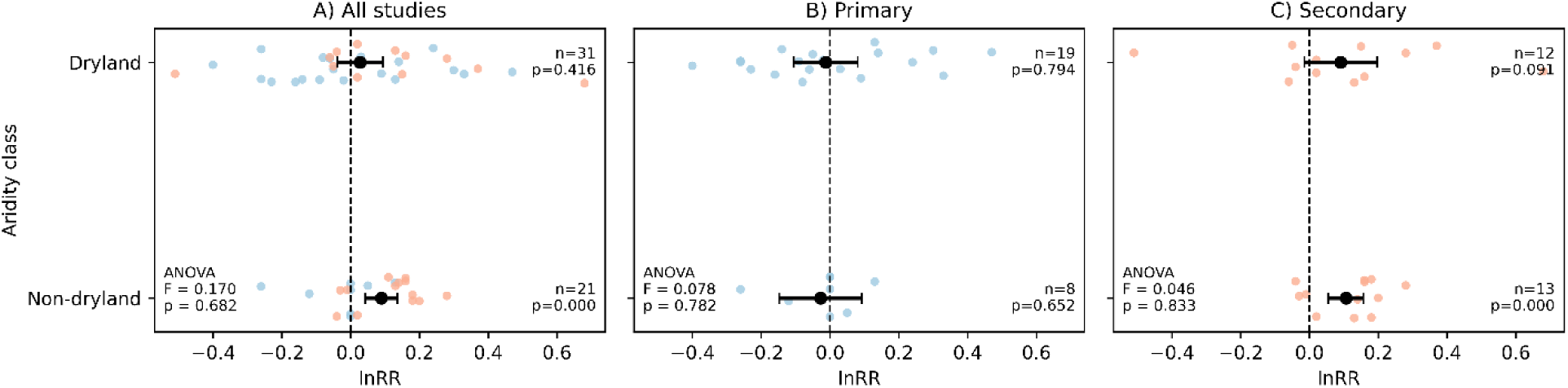
Soil organic carbon (SOC) effect size measured as the natural-log of the response ratio (lnRR) in response to adoption of alternative grazing practices. Results grouped as dryland and non-dryland classes, with a threshold of 0.65 for the aridity index. Colored points (blue = primary, red = secondary) show individual effect sizes and black symbols show pooled random-effects meta-analytic estimates with 95% confidence intervals. ANOVA results shown for comparison between aridity classes for each study quality category.

**Figure 4.**
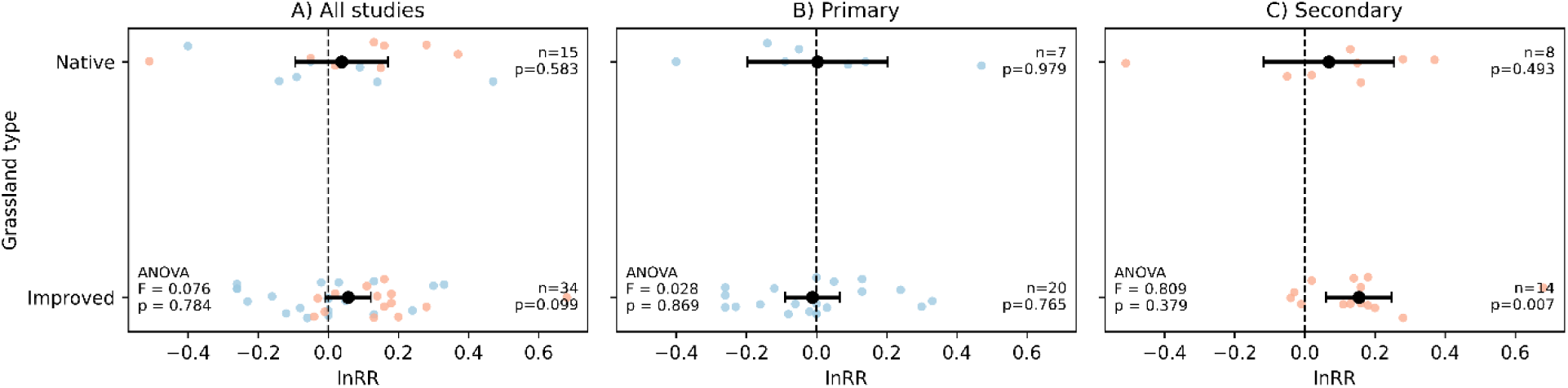
Soil organic carbon (SOC) effect size measured as the natural-log of the response ratio (lnRR) in response to adoption of alternative grazing practices. Results grouped by grassland type. Colored points (blue = primary, red = secondary) show individual effect sizes and black symbols show pooled random-effects meta-analytic estimates with 95% confidence intervals. ANOVA results shown for comparison between grassland type for each study quality category.

Given the near complete overlap between non-dryland climates and improved pastures, it is impossible to deduce whether it is the more mesic climate and/or the improved grass varieties and management (e.g. frequent resowing, liming, fertilization, irrigation) that lead to higher SOC, notwithstanding uncertainty in the representativeness of the controls calls into question the validity of this finding given that it is the secondary studies that are driving these trends.

Mean lnRR effect sizes, across all studies, were differentiated based on alternative grazing practice (Figure 5). Observations using adaptive management (i.e. animal movements and density fluidly managed based upon observations of forage availability and quality) had a small but significantly positive response (mean lnRR = 0.09, p = 0.017) that was also significantly greater than the mean of the prescribed grazing observations. This result is somewhat confounded by the fact that all but two observations from secondary studies used adaptive management. When limiting the data to primary studies, management type did not help explain the variability in SOC response (Figure 5). Unfortunately, due to lack of metadata describing key details of the grazing practices (i.e., stocking rates, frequency of movements, timing and duration of each grazing event), we could not explore management style beyond the reported categories of conventional grazing versus some form of investigator-defined alternative grazing management.

**Figure 5.**
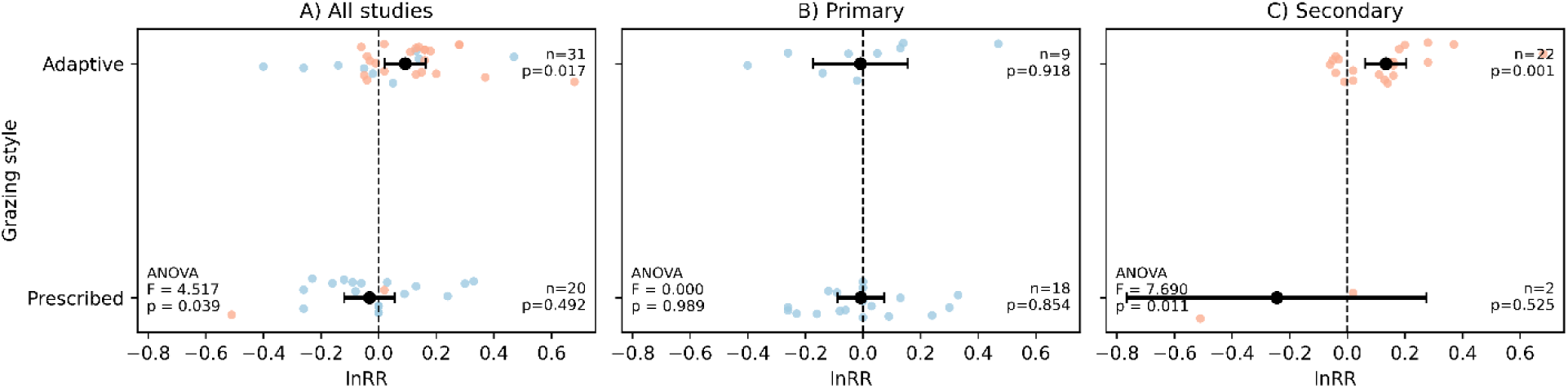
Soil organic carbon (SOC) effect size measured as the natural-log of the response ratio (lnRR) in response to adoption of alternative grazing practices. Results presented by grazing management style and presented by study quality category. Colored points (blue = primary, red = secondary) show individual effect sizes and black symbols show pooled random-effects meta-analytic estimates with 95% confidence intervals. ANOVA results shown for comparison between grazing styles.

## Discussion

When all retained data was pooled together, there was on average a slight positive SOC benefit to adopting an alternative grazing management system (mean lnRR = 0.05 (p = 0.06) and mean SOC sequestration rate = 0.29 tC ha^-1^ yr^-1^ (p = 0.08)). However, when we adhere to our strict inclusion criteria, slightly less than half the observations are excluded, and we find no SOC benefit upon adoption of alternative grazing management systems (mean lnRR = −0.02 (p = 0.21) and mean SOC sequestration rate = 0.03 tC ha^-1^ yr^-1^ (p = 0.90)). This finding suggests that the assertion of SOC benefits of grazing management has been predominantly informed by lower quality evidence. Reanalyzing previous meta-analyses (Supplement S5) using our systemic criteria suggests that earlier conclusions of large and significant SOC response in those studies, upon adoption of alternative grazing strategies, are not justified.

When only considering the primary studies, the data does not support any of our hypotheses around climate, grassland type or grazing style. Interestingly, the secondary studies, which comprise mostly adaptively managed livestock systems, do suggest that more mesic systems with improved grasses have a greater ability to sequester SOC than drier native grasslands. These findings are intriguing and point to the need for new concerted research efforts for two reasons: 1) we have underlying concerns about relying on one-off post-intervention measurements of paired sites, especially where the controls may not be representative of treatment conditions; and 2) aridity and grassland type are confounded with all improved grasslands falling into the non-dryland climate class.

How can scientists and practitioners work together to build a strong evidence base? An ideal grazing experiment would adopt a before-after-control-intervention (i.e. BACI) design implemented on a commercial scale and follow SOC changes for 5-10 or more years to provide strong causal evidence. Because this is logistically difficult to achieve, especially given typical short funding cycles, most investigations have taken a paired site approach with a one-time sampling campaign (i.e. a space-for-time substitution). Paired site studies can provide higher quality evidence when there is careful consideration of differences between treatment and control sites. In addition to the need to pair sites by climate, vegetation, soil type and topographic position, the underlying assumption of a paired site study is that the treatment and control sites were managed similarly for at least several decades prior to the implementation of the alternative grazing plan, and the control site continues the same management. Notably, we had 16 observations from four studies in the paired-site study design category for our primary studies, that met these criteria. However, it was much more common that this criterion was not met, with 34 studies either failing the criterion or simply not reporting the critical information needed for the criterion to be assessed. When they did report^25^, the information exemplified the challenge of finding representative controls. For example, in one pairing the alternative grazing site was converted from cropland 29 years prior to soil sampling whereas the continuous grazing control was converted 17 years prior. The difference in SOC stocks found in this pairing may simply be due to the extra 12 years under perennial pasture relative to cropland.

The controlled experimental studies had representative controls but likely because these studies were conducted at scales far smaller than the typical scale of management for commercial grazing operations^26^, with some plots as small as a few hundred square meters^27,28,29,30^, making it more logistically feasible to establish representative controls. Additionally, our finding of smaller effect sizes in controlled experiments than in observational studies overall runs counter to typical agronomic results where highly controlled plot-scale experiments have larger effect sizes than seen in farm-scale commercial settings^31^. It is difficult to know the scale-dependence of the SOC responses to grazing management, but it makes for a difficult case to say these small-scale studies were representative of real-world conditions^32^. However, we highlight that in our primary category we had an approximate balance between highly-controlled studies with higher internal validity and paired-site designs with higher external validity, and both designs estimate no strong effect of alternative grazing on SOC stocks (Table S4). Our findings should stand as a call to the research community that large-scale interventional studies are needed that go beyond paired-site designs to include repeated measurement of representative treatment and control ranches, such as BACI designs, if we want high quality, externally valid, evidence as to whether, and if so under what conditions, alternative grazing strategies have the ability to increase SOC stocks.

A common thread in both the primary literature and influential meta-analyses on the topic of SOC response to grazing management^13^ is lack of data transparency. Most of the studies only reported mean data, with 10 out of 23 neglecting to report variance metrics on individual treatments. Only one study provides the raw data behind the reported mean responses, and more than half did not include a data availability statement (see Supplemental Materials). Much has been written on this topic^33^ but it bears repeating that science needs to be replicable and reproducible to build confidence that recommendations will have desired outcomes, requiring open sharing of underlying data.

The lack of strong evidence of SOC benefits upon adoption of alternative grazing management leaves a void where proponents of these grazing strategies can continue to either cite lower quality studies or rely on anecdotal accounts of positive experiences. Notably, the secondary quality studies were cited on average three times as often per year as the primary studies (Table S6), suggesting that the secondary studies are having more influence on driving practitioner and investor opinions.

Given the rapidly growing interest in alternative grazing management practices as a NbCS, there is a critical need to establish a much more robust evidence base^34^. Such evidence should include controlled longitudinal experiments, using fit-for-purpose causal designs with protocols and tools that can quantify SOC change due to management at the scale of whole ranch management^35^. Such evidence will permit confident evaluation of whether or not investments in alternative grazing management will result in real climate benefits.

## Methods

### Literature search

Peer-reviewed investigations of the impact of switching from conventional continuous grazing to alternative grazing strategies on SOC were compiled using four different sources. Slightly different search strings were used for each source due to limitations on the number of Boolean terms.

Web of Science search string:

search for:((ALL=(soil)) AND ALL=("grazing management" OR "rotational grazing" OR "continuous grazing" or "adaptive management" OR "time controlled grazing" OR "deferred grazing" OR "cell grazing" OR "holistic management" OR "grazing systems" OR "grazing strategies")) AND ALL=("soil organic carbon" OR "soil carbon" OR "carbon sequestration" OR "soil organic matter" OR "C sequestration" OR "SOC" OR "SOM" OR "soil C" OR "OC" OR "TOC"OR "chemical characteristics")

Elsevier Science Direct search string:

searched: ALL=(soil) AND ALL=("grazing management" OR "rotational grazing" OR "continuous grazing") AND ALL=("soil organic carbon" OR "soil carbon" OR "carbon sequestration")

Google Scholar & Scopus (via Publish or Perish^36^) search strings:

search for: Soil organic carbon "rotational grazing" OR "continuous grazing" OR "adaptive management" OR "time controlled grazing" OR "deferred grazing" OR "cell grazing" OR "holistic management" OR "grazing systems" OR "grazing strategy” AND “carbon sequestration” OR “SOC” OR “C stocks”

These searches were conducted in June 2023. The completeness of our search was assessed by the successful retrieval of a short list of key studies including all of the papers used in the Byrnes et al.^35^ meta-analysis ^37,38,39,40,41,42,43,44,45,46,47,48^. Five additional primary publications were found in a meta-analysis of grazing impacts on SOC in Australia^14^ that were not included in our initial search, but none were found to pass our inclusion criteria.

### Filtering and Inclusion criteria

Abstracts from all returned studies were analyzed by asking the following three questions:

1. Is there a comparison of alternative (defined using any of our keywords above) and conventional (i.e. continuous or season-long) grazing strategy? Studies that only tracked SOC under alternative practice without a comparable continuous control baseline were excluded because of the difficulty in disentangling management from climate-induced changes in SOC^49^
2. Are SOC outcomes reported?
3. Is this primary research? Often multiple studies will use the same primary dataset in different ways such as for modeling or life cycle analyses. We wanted to be sure that the same field measurements weren’t being used more than once in our meta-analysis.

Papers were only retained if the answer to all three of these questions was yes.

After this filtering, the retained papers were read in detail to determine whether the study met our predefined inclusion criteria (Table 1). After reading the papers in detail, it was determined that several studies failed at least one of the initial questions – e.g. an alternative grazing management might have been mentioned in the abstract but upon reading the methods, there wasn’t an actual comparison with conventional grazing.

Knowledge of the method of carbon determination, especially in semi-arid systems where carbonates are often present, is important for determining the quality of the SOC data. We chose three years as the minimum study duration as SOC changes slowly and it is extremely unlikely that real differences can arise in treatments in less than three years. While topsoil (≤ 10 cm) contains greater SOC% relative to other depths, in grasslands the topsoil only accounts for a small fraction of the total SOC stocks and changes due to management in this layer may not be representative of changes in the entire profile. The question on significant external factors was included because the raw means in regional survey studies cannot be compared until potential confounding factors (climate, soil, topographic, etc.) that also influence soil carbon stocks and their rate of change are accounted for (see Allen et al.^36^ for a good example in the grazing-SOC literature). Accounting for these factors builds confidence that estimates of soil carbon change are due to the grazing management and not non-target causal variables like mean annual precipitation. Similarly, the representativeness of the control site in observational paired-site studies must satisfy expectations that treated and control sites are similar enough with regards other potential causal variables, such as soil texture and longer-term management history, to permit differences in soil carbon stocks to be attributed to grazing management. When such paired-site studies are conducted at a single point in time, as opposed to including pre-intervention data on soil carbon stocks that can be used to calculate differences due to treatment versus control of changes in time, appropriate matching of pairs by other potential causal variables becomes critical because SOC levels are unknown at the time of practice adoption, invoking the assumption that they were approximately equal at that time. That is, we have to be confident that the baselines (i.e. controls) for the sample of sites that are measured are representative enough in terms of environmental, soil, land use and historical management variables for them to allow confidence that estimated effects of grazing management are not due to non-random bias in observed and/or non-observed non-target causal variables.

Studies that met all these criteria are termed primary studies in all subsequent analyses. If a study met all the criteria except the final one on representativeness of the control site, the study was retained but flagged as secondary quality. These secondary studies were kept in the analysis because the majority of studies of alternative grazing management studies have this paired-site design.

### Data extraction

For all retained studies, the following information was extracted or at least attempted to be extracted: study year, experimental design, treatments, whether the alternative grazing strategy was adaptive or prescriptive (i.e. decisions on animal density and movements made based on observations of forage availability/quality vs. following a set plan regardless of seasonal conditions), controls used, land use history, duration of treatment on land prior to sampling, grassland type (native or improved) and species, methods for improving the pasture (e.g. fertilizer, plantings, etc.), size of treatment area/study scale, grazing animal type, stocking rate, tools used for soil sampling, maximum depth of sampling, soil sampling strategy, method for determining SOC content, carbonate removal information, SOC content and stock data, bulk density (BD) data, country, coordinates, mean annual precipitation (MAP), mean annual temperature (MAT), and world soil order. If climate data or soil order was not available it was extracted using location data from ClimateCharts.net^50^ and SoilGrids^51^. Aridity index (AI) was also extracted using location data from the Global-Aridity_ET0^52^.

A mean SOC concentration (wt%) was calculated for the 0-30 cm and 0 to maximum depth increments by taking the BD-weighted means for each horizon within the specified depth range. SOC stocks (tC ha^-1^) were either directly extracted from the study or calculated based on SOC concentration and BD and depth. No attempt was made to correct for equivalent soil mass.

We attempted to extract variance for each study. For studies where standard deviation (SD) or standard error (SE) were directly reported, we extracted these data, converting SE to SD by using the sample size. Ten of the 23 studies did not report SD or SE (and the underlying data was not available); instead, we used reported confidence intervals, least significant differences, or standard error of the mean from ANOVA testing to approximate a SD for each treatment. In several cases, we reduced SOC% and BD variance to coefficients of variation to estimate SD for the whole profile stocks. Where all of these steps failed, we simply assumed a coefficient of variation (CV) of 15% for SOC% and SOC stocks. Additionally, it was difficult to back out sample size for several of the studies, so we made our best guess based on the reported methods. The supplemental data file details how we estimated variance for each study.

All data from our synthesis, including the full list of studies retrieved from the literature search, the short list of 70 studies along with their assessment against our inclusion criteria, and the extracted data from the final list of studies are available online (https://doi.org/10.6084/m9.figshare.30660842). The locations of the final primary and secondary studies are shown in Figure 6.

**Figure 6.**
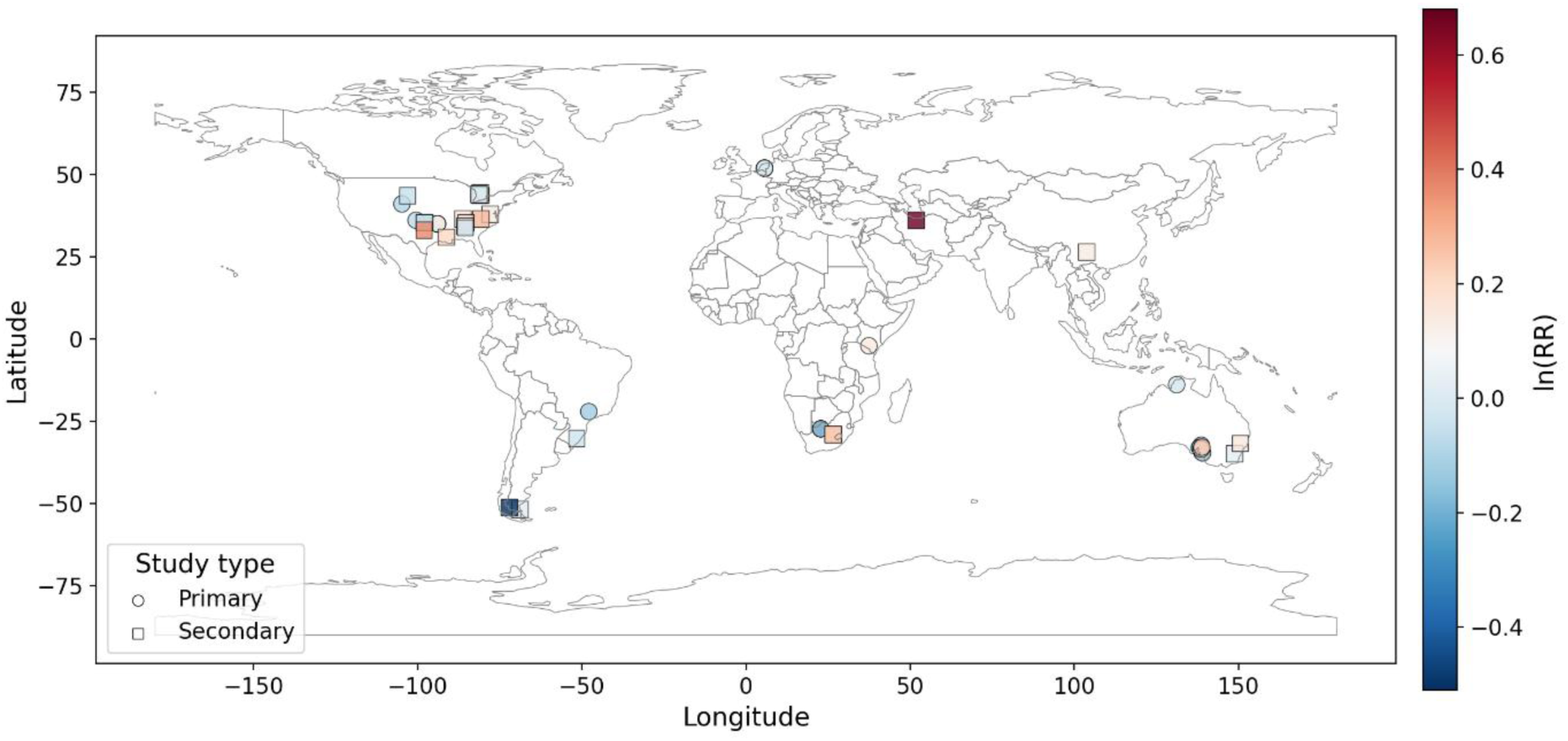
Locations of primary and secondary studies. Site locations colored by natural-log of the response ratio between alternative and conventional grazing practice.

### Data analysis

Differences between continuous and alternative grazing practices were summarized in two ways. First, an effect size measured as the natural log of the alternative grazing strategy SOC response divided by the continuous grazing SOC response (lnRR) was calculated for different comparisons (see below). Uncertainty in lnRR for each individual study comparison was estimated as standard error (SE_lnRR_) using the Taylor expansion method:

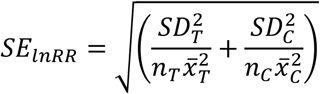

where SD, *n* and *x̅* are the standard deviation, sample size and mean SOC% or SOC stock of the treatment (*T*) and control (*c*) groups. Egger’s Regression^23^, where standardized effect sizes (lnRR/SE_lnRR_) are plotted against precision (1/SE_lnRR_), was then used to test for publication bias. Second, when all the necessary data was available, relative SOC sequestration rates (tC ha^-1^ yr^-1^) were also calculated for the 0-30 cm and 0-maximum depth ranges.

Initial analyses indicated that the effect size calculated from SOC concentration and SOC stocks were nearly identical (see Figure S1) and a one-way ANOVA indicated that there were no differences among studies that only measured SOC to < 30 cm, 30-50 and >50 cm (see Figure S2). These two findings allowed us to simplify our analyses by only reporting one effect size as the average of lnRR of SOC concentration and lnRR of SOC stocks, where both were reported, or just the lnRR of SOC concentration or stock where only one was reported, and by only reporting the results for 0-maximum depth range.

For the response-ratio analysis, effect sizes were calculated as the natural log response ratio (lnRR) for each observation. Sampling variances were derived as described above, and a random-effects meta-analysis^53,54^ was conducted to estimate the overall mean effect size and its 95% confidence interval. Individual observations were weighted based on the inverse of their variance. Because multiple observations were available from some studies, study ID was included as a random effect. Effect sizes are presented as pooled mean lnRR values with 95% confidence intervals and associated p-values (difference from zero).

For SOC sequestration rates, individual observation-level standard errors could not be calculated for most studies, so a conventional inverse-variance meta-analysis could not be performed. Instead, we used a study-clustered summary approach^55,56^. First, SOC sequestration values were averaged within each study so that each study contributed a single mean value.

The mean SOC sequestration across studies was then calculated, and uncertainty was estimated from the among-study variation in these study means. Standard errors, 95% confidence intervals, and p-values were therefore based on the distribution of study-level means rather than on observation-level sampling variances.

Results were analyzed as described above for the following categories: 1) study quality, 2) experimental design, 3) alternative grazing strategy, 4) aridity index and 5) rangeland type (natural versus introduced/improved grasses). For experimental design, the groups were limited to controlled experiment and paired site design because those two groups represented 94% of the data. The aridity index (AI) was binned into two categories: AI > 0.65 was considered ‘non-dryland’ and AI < 0.65 considered ‘dryland.’ For alternative grazing management, we sought to categorize studies into adaptive versus prescribed management where prescribed means a set grazing schedule was adhered to and adaptive means the grazing schedule was adjusted based on observations of actual forage consumption and condition. We would have liked to try to define alternative grazing strategies based on quantitative measures of stocking rate, frequency and duration but these data were not provided in > 25% of the publications (see data supplement). For pasture type, the groups were limited to improved or native pasture. One-way analysis of variance (ANOVA) tests were conducted to test for significant differences between classes in each categorical comparison. ANOVA assumptions were tested for each test. Across all tests, when data were pooled into all studies, assumptions of normality, homogeneity of variance and independence of observations were met. When testing within the primary or secondary study categories, normality and homogeneity assumptions were met but the independence assumption isn’t as robust given that a small number of studies contributed an outsized number of observations. All analyses and plotting were done in Python using the Pandas^57^, Seaborn^58^, Matplotlib^59^ and SciPy^60^ libraries.

## Data Availability

The full list of papers, our inclusion/exclusion criteria applied to each paper, and the final extracted data that were used to generate the findings in this study are available at figshare with the following identifier: https://doi.org/10.6084/m9.figshare.30660842

## Acknowledgements

Support for Woodwell Climate to conduct this work came from National Fish and Wildlife Foundation, the Mighty Arrow Foundation and the JM Kaplan Fund. This work was also supported by the Environmental Defense Fund with awards from King Philanthropies and Arcadia, a charitable fund of Lisbet Rausing and Peter Baldwin. MAB was involved with the work through the Yale Applied Science Synthesis Program, a joint initiative of The Forest School (at the Yale School of the Environment) and the Yale Center for Natural Carbon Capture, through support from Mary Anne Nyburg Baker and G. Leonard Baker, Jr.

## Author Contributions

Jonathan Sanderman: Conceptualization (lead), formal analysis (lead), methodology, writing – original draft

Colleen Partida: Data curation (lead), investigation, writing – original draft, review and editing Yushu Xia: Conceptualization, writing - review and editing

Jocelyn M. Lavallee: Conceptualization, writing - review and editing Mark A. Bradford: Writing - review and editing

## Supplemental Information

### S1. Publication bias

To test for the potential of publication bias, the lnRR values and their associated standard errors were used in an Egger’s Regression and visualized as Funnel plots for all studies, primary studies only and secondary studies only (Figure S1). None of the slopes of these regressions were significantly different from zero suggesting that there was no publication bias in the final selection of studies.

**Figure S1.**
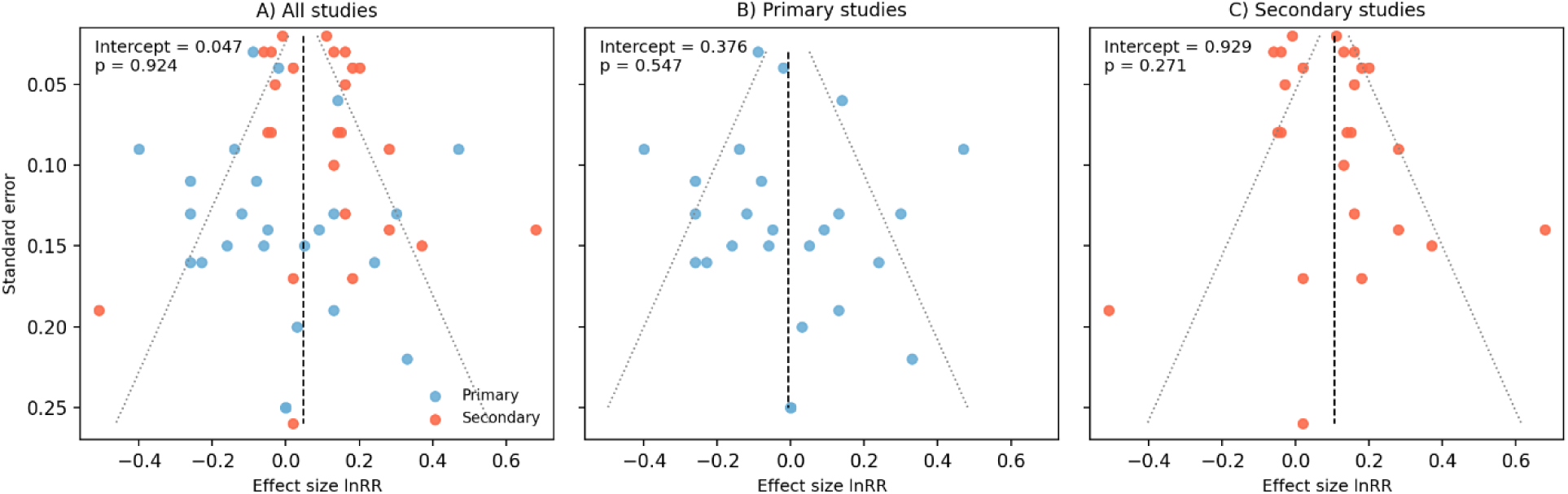
Funnel plots of the natural-log of the response ratio (lnRR) for soil organic carbon response to a shift in grazing management. Results from Egger’s regression are given in upper left corner of each panel.

### S2. Reporting SOC concentration or stocks did not matter

The lnRR results from SOC concentration (wt%) and SOC stocks were nearly identical (Figure S2). As such, in subsequent analyses the lnRR was calculated as the average of the lnRR for SOC% and SOC stocks. This allowed the maximum number of comparisons to be included in statistical tests because not all studies reported SOC% and stocks.

**Figure S2.**
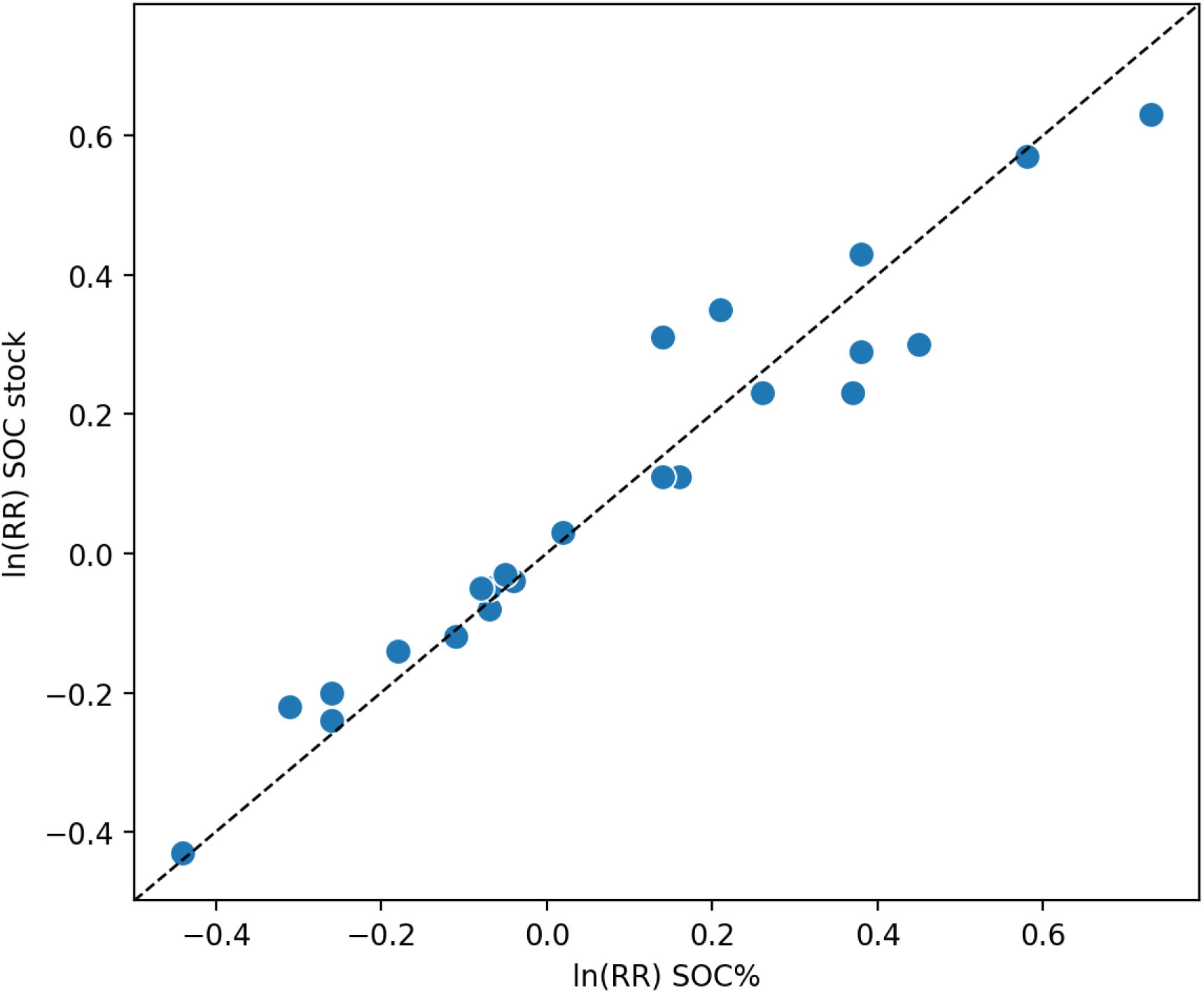
Comparison of natural-log of the response ratio (lnRR calculated for SOC concentration (wt%) and for SOC stocks (tC ha^-1^) where data was available to calculate lnRR both ways (n = 22) indicates that there is no systematic difference in lnRR allowing pooling of data to maximize sample size in the meta-analysis.

### S3. No clear effect of soil measurement depth

Perhaps due to our inclusion criteria that sampling depth needed to be greater than 10 cm, the mean sampling depth was 50 cm and, importantly, there was no difference in results between studies measuring to less than 30 cm, those that measured to between 30-50 and those that measured greater than 50 cm (Figure S3). Single factor ANOVA suggested no difference between these three groups (ANOVA F = 0.589, p = 0.559). Results for SOC sequestration rates were similar (data not shown). This result gave us confidence in grouping all depths together for the main statistical analyses.

**Figure S3.**
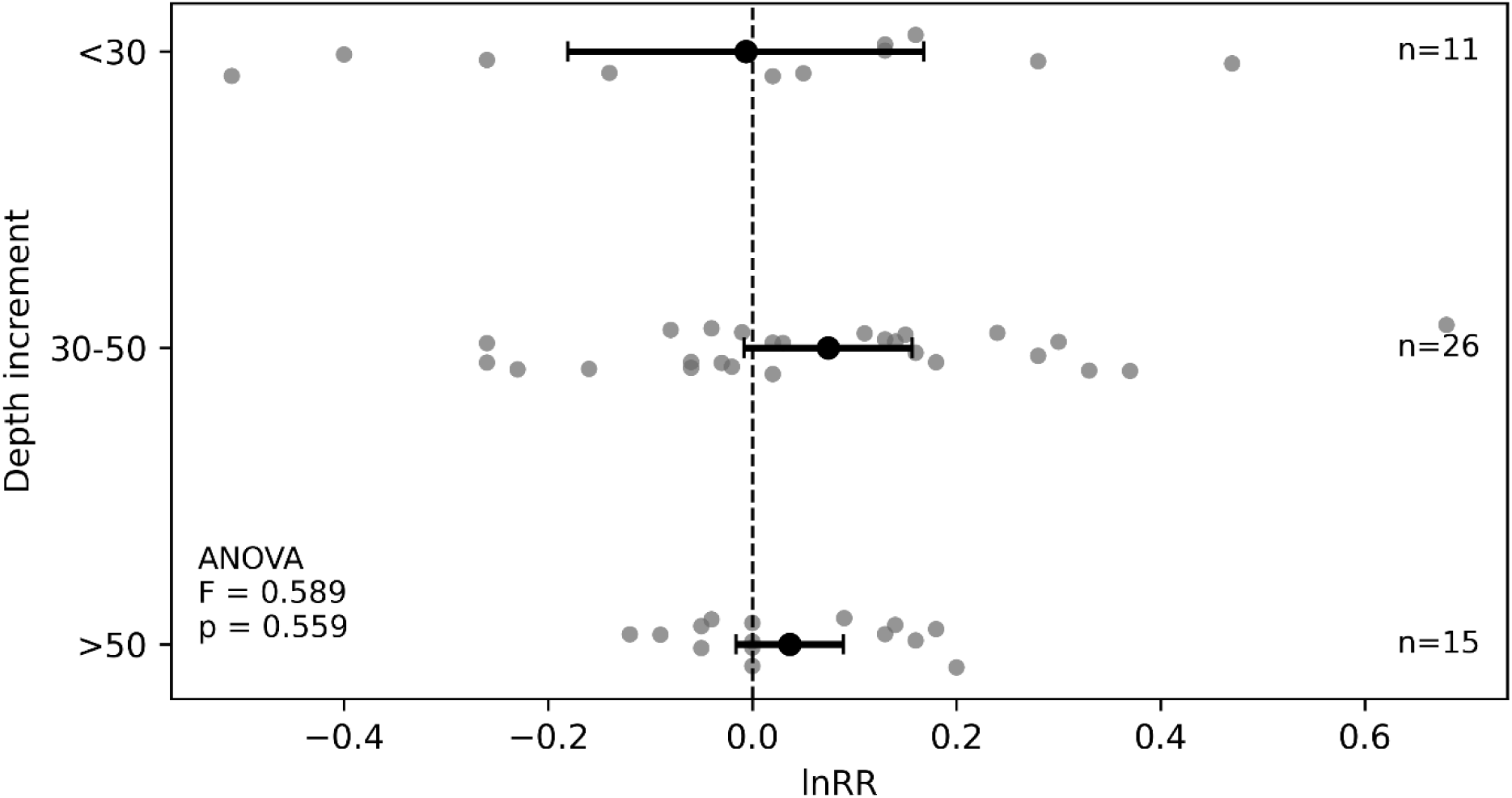
Natural-log of the response ratio (lnRR) for soil organic carbon grouped by total sampling depth. Points show individual effect sizes and black symbols show pooled random-effects meta-analytic estimates with 95% confidence intervals. ANOVA results shown for comparison between depth groups.

### S4. Paired site studies dominate positive responses

There were four study designs found in our final paper list: controlled experiments where an area was divided into multiple control and intervention plots, unreplicated experiments where an area was divided into just one replicate of each control and intervention, paired site designs where two “paired” fields under contrasting management were compared, and survey designs where a number of ranches with varying management practices across a region were measured. Only one study adopted the survey design and only two were unreplicated experiments. These three observations were excluded from this analysis of study design. Figure S4 indicates that nearly all the positive SOC responses were associated with paired site studies rather than controlled experiments, but these were primarily the same studies that dominated the secondary study class based on the inclusion/exclusion criteria. Pooled mean lnRR was no different than zero for the controlled experiments (p = 0.87) while the paired sites were greater than zero with a p value = 0.09; however, one-way ANOVA results (F = 1.112, p = 0.297) suggest these two categories are not that different from each other. This result is driven by the large dispersion in each dataset.

**Figure S4.**
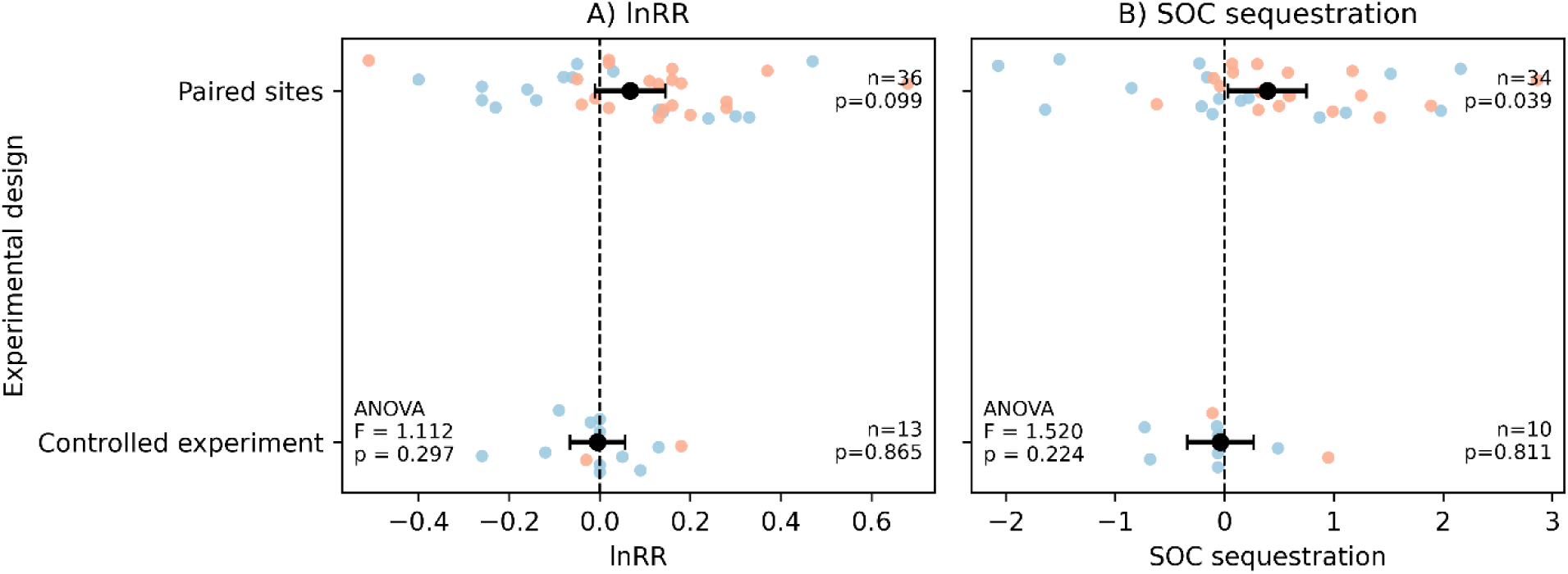
Soil organic carbon (SOC) effect size measured as the natural-log of the response ratio (lnRR) and annual sequestration rate in response to adoption of alternative grazing practices grouped by study design. Colored points (blue = primary, red = secondary) show individual effect sizes and black symbols show pooled random-effects meta-analytic estimates with 95% confidence intervals. Results combined for primary and secondary studies. ANOVA results shown for comparison between experimental designs.

There were no differences in SOC response between study types when only considering primary studies (Table S4). The difference between study designs for the secondary studies could not be assessed because this category was dominated by paired site studies; however, within only the paired-site secondary studies, lnRR and SOC sequestration rate were both significantly different from zero (Table S4).

**Table S4.**
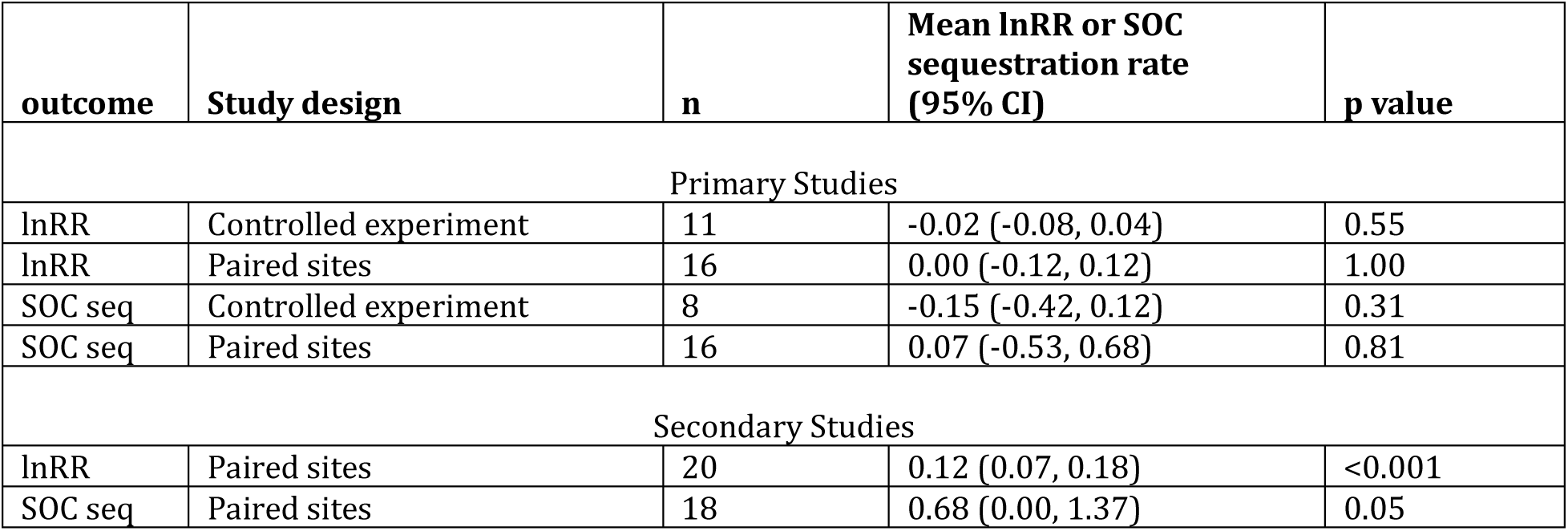
Pooled effect sizes when the primary studies are separated into experimental design and secondary studies are limited to only paired site designs. The sample size was too low for secondary controlled studies.

### S5. Reconsidering existing literature

Given that Byrnes et al.^4^ has been one of the main studies cited when supporting the notion that the adoption of alternative grazing strategies leads to SOC sequestration, we tried to determine why our conclusions varied so much from the conclusions in that study. Byrnes et al.^4^ supply a list of studies that were used in estimating effect sizes but not the detailed data themselves making it difficult to reproduce their results. We re-extracted the data and attempted to recreate their analysis but couldn’t replicate their findings (Table S8). Byrnes et al.^4^ report a mean effect size of 0.25 (95% CI: 0.10 - 0.41) for 44 comparisons of alternative and conventional grazing strategies but we could only reconstruct a mean effect size of 0.15 for the same number of comparisons for the Byrnes et al.^4^ dataset. Part of this difference may be due to different choices in how to handle variance and multiple comparisons in an individual study.

More importantly, most of the data sources used by Byrnes et al.^4^ did not meet our inclusion criteria (Table S6), highlighting the need to consider the quality of the studies being assembled for quantitative syntheses if one is to have confidence in the average effect across studies^5,6^. In fact, 7 of the 12 studies listed were omitted from our meta-analysis for failing at least one of our inclusion criteria or repeating data already included in another study. Another 4 studies made our secondary list and only one study was on our primary study list. To approximate the same sample size for each study as reported in Byrnes et al.^4^, it appears that each soil depth increment must have been included as an independent observation (i.e. if a study measured 0-5, 5-15 and 5-30 cm there were three data points used instead of just one data point for 0-30 cm).

A notable inclusion by Byrnes et al.^4^, which we excluded because it failed exclusion criteria #5 (Table 1), was Allen et al.^7^. In this regional survey in Queensland, Australia, the authors measured much higher SOC stocks under alternative grazing strategies than under conventional grazing. However, after the authors of the original paper conducted a statistical analysis that removed the confounding effect of climatic differences among sites (all the alternative grazing sites were in wetter regions of the state), they concluded that pastures under alternative grazing strategies had less SOC than conventionally grazed pastures. Byrnes et al.^4^ apparently extracted and included the raw data without consideration of the need to account for confounding in the study design.

There are a few high-profile papers that examined a shift from cropland to alternative grazing management of grasses^8,9^ which are often incorrectly used to support the notion that alternative grazing strategies can have large SOC benefits. Both studies focused on chronosequences of improved pastures seeded into former cropland in the SE of the United States. Large gains in SOC typically occur when croplands are converted to pastures regardless of grazing management strategies^10^. A third highly-cited paper concluding large sequestration rates under an adaptive grazing plan^11^ was also excluded from our study because there was no comparison to continuous grazing, making attribution of any change over time in SOC to the grazing management strategy difficult.

**Table S5.** Comparison of our attempt to reconstruct results from Byrnes et al.^4^ and our extraction (this study) of the same data sources. Studies were classified using our inclusion criteria (see methods).

| Study quality | <u>Byrnes reconstruction</u> |  |  | <u>this study</u> |  |  |
| --- | --- | --- | --- | --- | --- | --- |
|  | N | Mean | 95% CI | n | mean | 95% CI |
| Primary | 9 | -0.01 | (-0.31, 0.29) | 3 | 0.01 | (-0.57, 0.59) |
| Secondary | 17 | 0.22 | (0.15, 0.29) | 6 | 0.24 | (0.13, 0.35) |
| Omitted | 18 | 0.17 | (0.05, 0.28) |  |  |  |
| **All** | **44** | **0.15** | **(0.07, 0.23)** | **9** | **0.16** | **(-0.04, 0.36)** |

### S6. Citation frequency

We used Google Scholar to capture the number of times each study was cited as of June 2025 because Google Scholar picks up both peer-reviewed and grey literature articles and reports, making it a more inclusive measure of the impact of a particular study. The 10 primary studies were cited 225 times, while the 13 secondary studies were cited a total of 1508 times with, on average, 3 times as many citations per year (Table S9).

**Table S6.** Google Scholar citation rate for studies group by study quality.

| **Study quality** | **<u>Sum of citations</u>** | **<u>Average citations per study</u>** | **<u>Average citations per study per year</u>** |
| --- | --- | --- | --- |
| Primary | 287 | 29 | 4.3 |
| Secondary | 1508 | 116 | 12 |
| **All** | **1795** | **78** | **8.7** |

